# Top-heavy trophic structure within benthic viral dark matter

**DOI:** 10.1101/2022.10.01.510451

**Authors:** Ethan C. Cissell, Sophie J. McCoy

**Author notes:** Address correspondence to: Ethan C. Cissell, **Email:**. **Author Contributions:** E.C. Cissell conceived of and designed the study with significant contributions from S.J. McCoy. E.C. Cissell led all data collection in the field, and all in-situ data analyses and bioinformatics. S.J. McCoy significantly contributed to in-situ data analysis. E.C. Cissell drafted the original manuscript with significant contributions from S.J. McCoy. Both authors contributed significantly to manuscript revision. **Competing Interest Statement:** The authors declare they have no known competing financial interests or personal relationships that did or could have appeared to influence the work reported in this paper.

## Abstract

Viruses exert considerable influence on microbial population dynamics and community structure, with cascading effects on ecosystem-scale biogeochemical cycling and functional trajectories. Creating broadly generalizable theory on viral trophic ecology requires further inquiry into historically unexplored microbial systems that currently lack empirically demonstrated patterns in viral infectivity, such as structurally complex benthic communities. This becomes increasingly relevant considering recently proposed revisions to the fundamental mechanisms that modulate the strength and direction viral trophic linkages. Here, we employed deep longitudinal multiomic sequencing to characterize the viral assemblage (including *ss*DNA, *ds*DNA, and *ds*RNA viruses) and profile lineage-specific host-virus interactions within benthic cyanobacterial mats sampled from Bonaire, Caribbean Netherlands, over a complete diel time-series, and reconstruct patterns in intra-mat trophic structure. We recovered 11,020 unique viral populations spanning at least 10 viral families across the orders Caudovirales, Petitvirales, and Mindivirales, with evidence for extensive genomic novelty from reference and environmental viral sequences. Analysis of coverage ratios of viral sequences and computationally predicted hosts spanning 15 phyla and 21 classes revealed virus:host abundance and activity ratios consistently exceeding 1:1, with overall power-law scaling indicating an increasingly top-heavy intra-mat trophic structure with significant top-down pressure. Diel activity of cyanophages showed clear temporal patterns that seem to follow host physiological condition. These data generate important hypotheses concerning taxon-dependent variation in the relative contribution of top-down vs. bottom-up forcing in driving mat community dynamics, and establish a useful database of viral sequences from this previously unexplored system toward the generation of generalizable trans-system theory on viral trophic ecology.

**SIGNIFICANCE STATEMENT:** Recent advances in viral ecological theory suggest a better understanding of system-specific viral ecology is needed from diverse environments to create generalizable theory that accurately predicts patterns of trophic interaction strengths across systems, especially in the Anthropocene. This study characterized viral-host trophic structure within coral reef benthic cyanobacterial mats - a globally proliferating cause and consequence of coral reef degradation - using paired multiomic sequencing. Recovered viral sequences displayed remarkable genomic novelty from other well-characterized viruses and spanned diverse viral taxa. Unexpectedly, lineage-resolved trophic linkages displayed a strongly active top-heavy trophic structure, suggesting extensive top-down forcing. These results highlight the context-dependence of viral trophic interaction strengths and suggest that viruses strongly influence reef cyanobacterial mat and reef ecosystem functional trajectories.

## INTRODUCTION

Viruses ubiquitously exert considerable influence on cellular microorganisms across scales of biological and ecological organization. At the cellular level, both viral lytic and lysogenic interactions modify cellular physiologic potential (1–3) and transcriptional regulation pathways (4), alongside mediating cellular susceptibility to top-down control from both homo- and heterotypic viral infection (5–8) and bacterivorous zooplankton (9). Transfer of exogenous genes into hosts and antagonistic coevolutionary dynamics promote extensive genomic diversification and steer population evolutionary trajectories (negative frequency-dependent selection; 10–12), alongside significant top-down control on population abundance via the outcome of viral lysis (13, 14). Top-down control on population size can promote coexistence among competing cellular microorganisms by preventing competitive exclusion by competitive dominants, contributing to patterns in microbial community structure and successional dynamics (5, 15, 16). These direct and indirect influence of viruses on microbial hosts in-turn scale to ecosystem biogeochemical cycling, environmental stoichiometry, and functional trajectory (17). For example, viral predation is thought to modulate planktonic marine carbon cycling, releasing approximately 10 billion tons of microbe-bound organic carbon into the Dissolved Organic Carbon (DOC) pool every day, and promoting retention of carbon within the basal trophic levels of the microbial loop (viral shunt) (18, 19). Understanding viral interactions with cellular microorganisms, then, fundamentally underlies an understanding of patterns in microbial diversity (both micro- and macro-), and is integral for scaling up microbial physiology, and population and community dynamics, to broader ecosystem processes (20).

Recent advances toward better *in-silico* methods of viral recovery from shotgun metagenomic and metatranscriptomic datasets have enabled the exploration of viral interaction dynamics across more diverse systems, and with greater resolution (21–23). Traditional paradigms in viral ecology are largely derived from few, structurally simple model systems (20, 24). However, emerging evidence from diverse structurally-complex systems challenges the generality of paradigms in viral ecology, especially in the directionality and scales of dynamism in both the direction and strength of viral trophic interactions (9, 25–27). Overall, these revisions from contemporary viral ecological theory suggest that current models linking viral interactions to community- and ecosystem-scale processes are likely incomplete, motivating a reinvigoration of basic research into viral ecology. Inquiry into historically unexplored microbial systems that currently lack empirically demonstrated patterns in viral infectivity are especially critical toward the creation of broadly generalizable theory and quantitative frameworks scaling viral trophic interactions to microbial community ecology and ecosystem chemistry (28–30).

Harmful algal blooms are an ideal system in which to investigate viral ecology, with motivation from both fundamental and applied perspectives. Planktonic cyanobacterial and eukaryotic algal blooms receive significant research and public attention. Yet, benthic blooms remain understudied, despite impacting coral reefs, which are some of the most threatened ecosystems globally (31). Benthic cyanobacterial mats are increasing in abundance on coral reefs worldwide (32) because of local and global stressors that are deleterious to the health of important reef building taxa (33–37). Cyanobacterial mats are taxonomically and functionally complex communities (38) that can dramatically alter reef nitrogen budgets via nitrogen fixation (39–41), and reef carbon budgets via fixation of inorganic carbon and subsequent release of dissolved organic carbon (42). These shifts in reef carbon chemistry can alter systemic microbial energetic budgets, and favor the proliferation and stability of pathogenic taxa (43), potentially including pathogenic taxa involved in coral disease (44). Horizontally-spreading reef cyanobacterial mat carpets are subject to dynamic predation pressure from generalist reef fishes and specialist invertebrate mesograzers (45, 46), with substantial geographic and morphotypic-dependence in predation risk from macro- and mesopredators (32, 47, 48). Viral interactions with coral reef benthic cyanobacterial mats have only recently been explored (38, 49, 50), and viral ecology in cyanobacterial mats remains generally relatively unknown.

Viral impact on microbial mat systems, including coral reef cyanobacterial mats, is not yet readily predictable without system-specific characterization (51, 52). The directionality of trophic interactions among viruses and their hosts may experience fine-scale dynamism dependent on density, host physiological state, and host trophic membership (27, 53), and thus may not be generalizable across time. Furthermore, living as a biofilm is generally known to confer some protection from viral predation (54), especially in spatially structured (generally biologically mediated) biofilm communities (25). This may presuppose the prediction of limited viral influence in coral reef cyanobacterial via similar mechanisms. However, viruses (evidence from bacteriophages) are also shown to subvert biofilm structural defense by exhibiting subdiffusive motion using mucin present in the extracellular polysaccharide matrix of biofilms to increase host encounter rates and overall infectivity in biofilms (55, 56). A more complete understanding of predation risk from viruses and the influence of viral trophic interactions on population mortality in cyanobacterial mats is critical toward both the generation of more generalizable theory, and for a better understanding of the processes governing cyanobacterial mat dynamics on reefs.

Here, we profiled lineage-specific host-virus interactions with ssDNA, dsDNA, and RNA viruses and reconstructed broad patterns in intra-mat tropic structure within cyanobacterial mats from Bonaire, Caribbean Netherlands, across a complete diel cycle using deep longitudinal multiomic sequencing (paired metagenomes and metatranscriptomes). Consistent with other high-density high growth-rate microbial systems (9, 26, 57), we hypothesized that viral predation (lytic interactions) would not be strongly evident in cyanobacterial mats.

## RESULTS

Four spatially distinct coral reef benthic cyanobacterial mats (Table S1) from Bonaire were longitudinally sampled at 5 evenly spaced time points across a complete diel cycle, and subject to paired shotgun metagenomic and metatranscriptomic sequencing, generating a total of 4,607,785,034 high-quality filtered reads (Table S2 & S3). *De-novo* coassembly of these reads was performed to reconstruct mat metagenomes and metatranscriptomes (Table S4 & S5), which were subsequently screened to isolate the cyanobacterial mat virome.

### Overview of viral community structure

We used combined profile Hidden Markov Models (HMM) and *BLASTp* with non-reference based HMM searches to extract viral signatures from coassembled metagenomes and metatranscriptomes, identifying a total of 13,026 manually curated viral sequences dereplicated at 95% identity within mat samples (approximating viral populations; vOTUs; 11,020 non-redundant vOTUs dereplicated across all samples; vMAT database). Recovered viral sequences included dsDNA, ssDNA, and dsRNA viruses, but primarily dsDNA viral sequences were recovered (Fig 1). A genomic protein-sharing network was constructed among viral sequences reconstructed in this study and viral sequences present in RefSeq_88_, and integrated with taxonomic predictions from trained convolutional-neural networks against the Caudovirales database to assign putative viral taxonomy, and explore viral diversity from reference viral sequences. Identified vOTUs could be assigned to 10 unique viral families, primarily belonging to the order Caudovirales (Fig. 1). Unsurprisingly, few viral sequences recovered in the vMAT database shared significant numbers of gene regions to viral sequences found in the RefSeq database, highlighting the highly novel diversity of viral sequences recovered from this system (Fig. 1b). Viral sequences assigned to the *Podoviridae* from convolutional-neural network analysis and those that shared edges (significant gene overlap) in the gene-sharing network with RefSeq *Podovirdae* sequence nodes displayed remarkably low edge betweenness centrality, suggesting that these sequences possess highly unique or divergent gene regions from other viral families (Fig. 1a).

**Figure 1.**
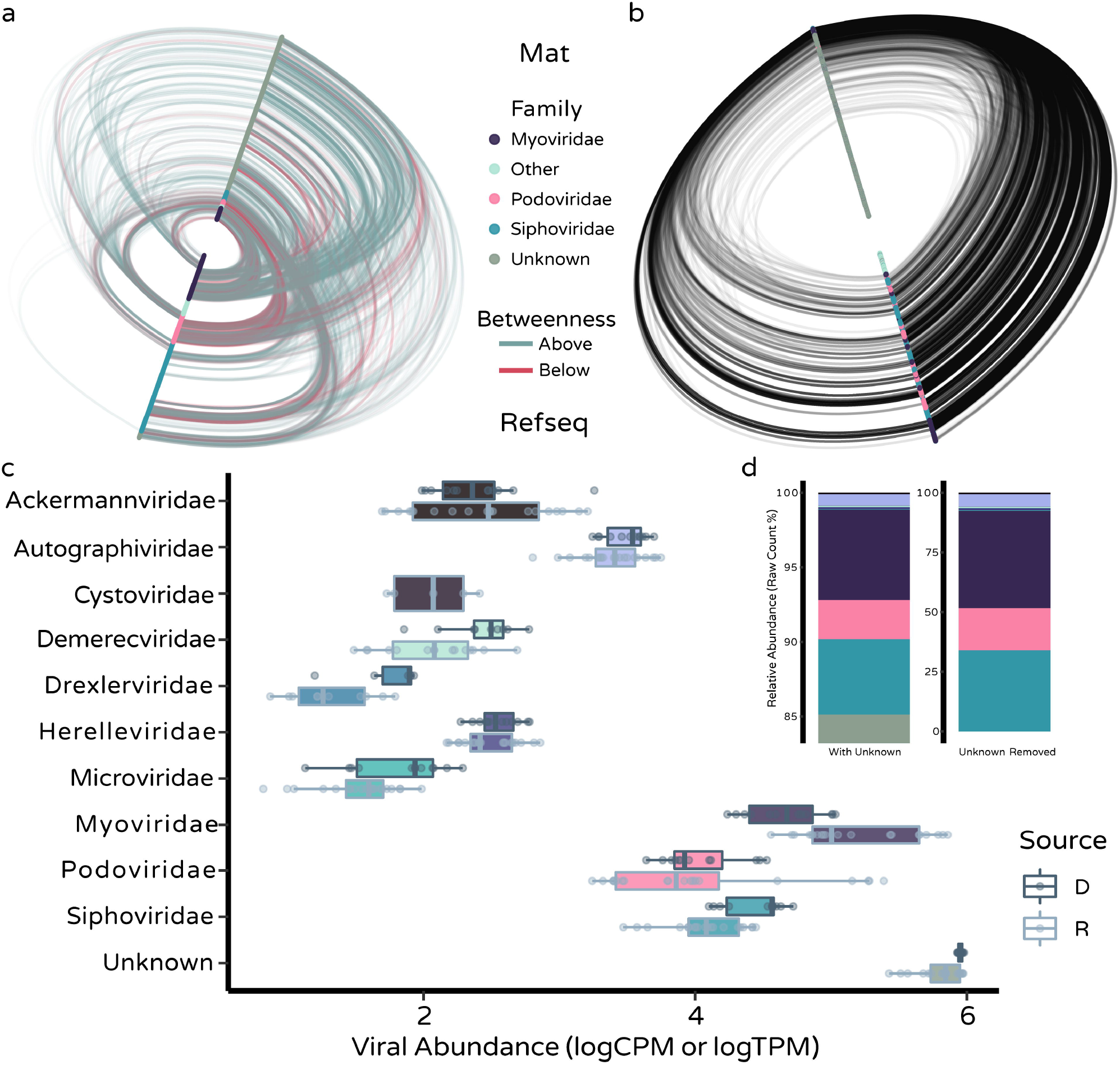
Cyanobacterial mat virome diversity and overview of abundances. Hive plot visualizations of gene-sharing network created among viral sequences recovered in this study and reference viral sequences (RefSeq_88_). Edges connect nodes that share a significant number of genes. Nodes are grouped on axes by dataset (Mat – top axis; RefSeq – bottom axis). Nodes are ordered along axes either by a, viral taxonomic (family) affiliation, or b, node betweenness centrality (high-low:outside-inside). In panel a, edges are colored by edge betweenness relative to median (above or below median edge betweenness). c, Boxplots of log transformed abundances (either CPM or TPM for metagenomic or metatranscriptomic read sets, respectively) across viral families. Box color denotes read-set source (D=DNA, R=RNA). d, Stacked barplots of proportional abundances of total number of viral sequences recovered from each viral family (colors from [c]) either including (left) or removing (right) viruses with unknown taxonomy.

Viral abundance and absolute sequence counts in metagenomes and metatranscriptomes was dominated by viral sequences that could not be placed within a resolved taxonomic lineage, followed by members of the *Myoviridae, Siphoviridae*, and *Podoviridae* (Fig. 1c, d). Viral sequences assigned to the *Cystoviridae* were also abundant in mat metatranscriptomes, suggesting RNA viruses are important members of cyanobacterial mat communities. Overview of viral sequences, including taxonomic assignments, are provided in Dataset S1.

### Cyanobacterial mat virome harbors substantial diversity unique from other environments

An additional genomic-based protein-sharing network was constructed among the viruses reconstructed from this study (vMAT), the viruses reconstructed from the Global Ocean Virome 2.0 (GOV2), and the viruses reconstructed from permafrost soil (vFROST) to contextualize the diversity uncovered in this dataset with other relevant publicly available environmental viral databases (Fig. 2a). Viruses recovered from the cyanobacterial mats sampled here displayed remarkable diversity from the sequences in these previously characterized environmental viral databases. Out of a total of 14,158 unique clusters (approximating genus-level relatedness), only 11 were shared among all datasets (Fig. 2b). As expected, viruses recovered from marine cyanobacterial mats (vMAT) shared fewer proteins and formed fewer unique clusters with permafrost viruses than with pelagic marine viruses (Fig. 2), likely owing to strong differences in abiotic filters between permafrost and benthic marine environments. The broad coverage of edge space across the diversity of the GOV2.0 dataset nodes suggests a broad commonality of gene regions across viruses in this dataset, with restricted distribution among viral sequences isolated from cyanobacterial mats, further highlighting the immense genetic diversity found in this system. More viral sequences were classified as singletons (n = 6,401) than those that clustered (n = 4,177), and the majority of those that clustered formed clusters exclusively with other cyanobacterial mat viral sequences (Fig. 2b). Taken together, these data reveal a high degree of endemism in mat community viruses, distinct even from viruses in surrounding seawater.

**Figure 2.**
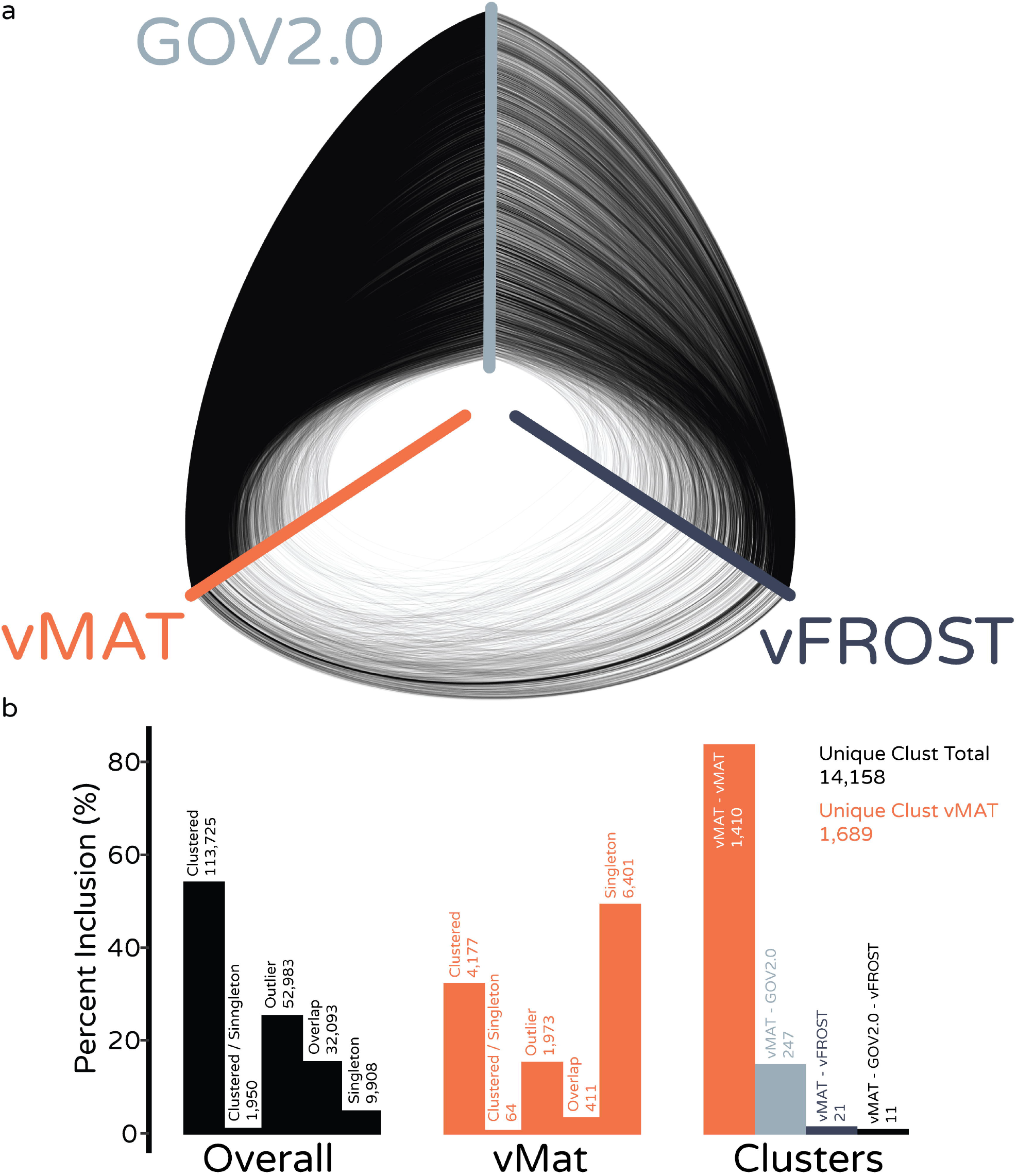
The cyanobacterial mat virome is unique from other relevant environmental viromes. a, Hive plot of gene-sharing network constructed among viruses reconstructed from cyanobacterial mats in this study (vMAT), viruses reconstructed from the Global Ocean Virome (GOV2.0) and viruses reconstructed from across a permafrost thaw gradient (vFROST). Edges connect nodes that share a significant number of genes, but do not necessarily encode clusters. Nodes are ordered along axes in order of decreasing node degree centrality (from exterior to interior). Nodes are grouped on distinct axes by dataset. b, Summary bar plots of clustering information from above (a) visualized gene-sharing network showing percent inclusion of nodes in each category of clustering status. Also given is raw node count above each bar, as well as total number of unique clusters among all nodes, and unique clusters including vMAT viruses.

However, there was an effect of viral sequence length on clustering status for vMAT viruses (PWRST, all p < 0.05; Fig. S1), which suggests genome incompleteness may, in-part, underly some patterns of purported extreme novelty within singletons. However, numerous long sequences were also classified as singletons, indicating genome incompleteness does not fully explain patterns in viral genomic novelty in the vMAT database.

### Viruses infect a broad host range in cyanobacterial mats

Multiple computational prognostic strategies for predicting host-virus infection relationships were integrated and hierarchically coalesced to reconstruct lineage-resolved host:virus infection pairs among viral sequences reconstructed in this study, and MAGs assembled in (Cissell and McCoy *in review*). Virus-host pairings were successfully assigned for 5,539 vOTUs (42.5% of all identified contigs) spanning 9 viral families, 14 bacterial phyla, 1 phylum of Archaea, and 21 classes (Fig. S2) of bacteria and Archaea (excluding 1 unresolved [class level] Proteobacteria linkage). These majority of these pairings were from vOTUs that could not be assigned to family level taxonomy (n = 4,734 linkages). To explore infection pairings among those viral sequences for which taxonomy could be predicted, infection data were subset to exclude those pairings without viral taxonomic assignment (Fig. 3). Of those pairings for which viral taxonomy was predicted, the bacterial phylum Proteobacteria numerically had the most predicted host viral linkages, with 315 pairings spanning 7 viral families (Fig. 3). Both Cyanobacteria and Bacteroidota were computationally predicted to be infected by members of 8 viral families, with infections dominated by the *Myoviridae* in both phyla (94 and 100 for Cyanobacteria and Bacteroidota, respectively; Fig. 3).

**Figure 3.**
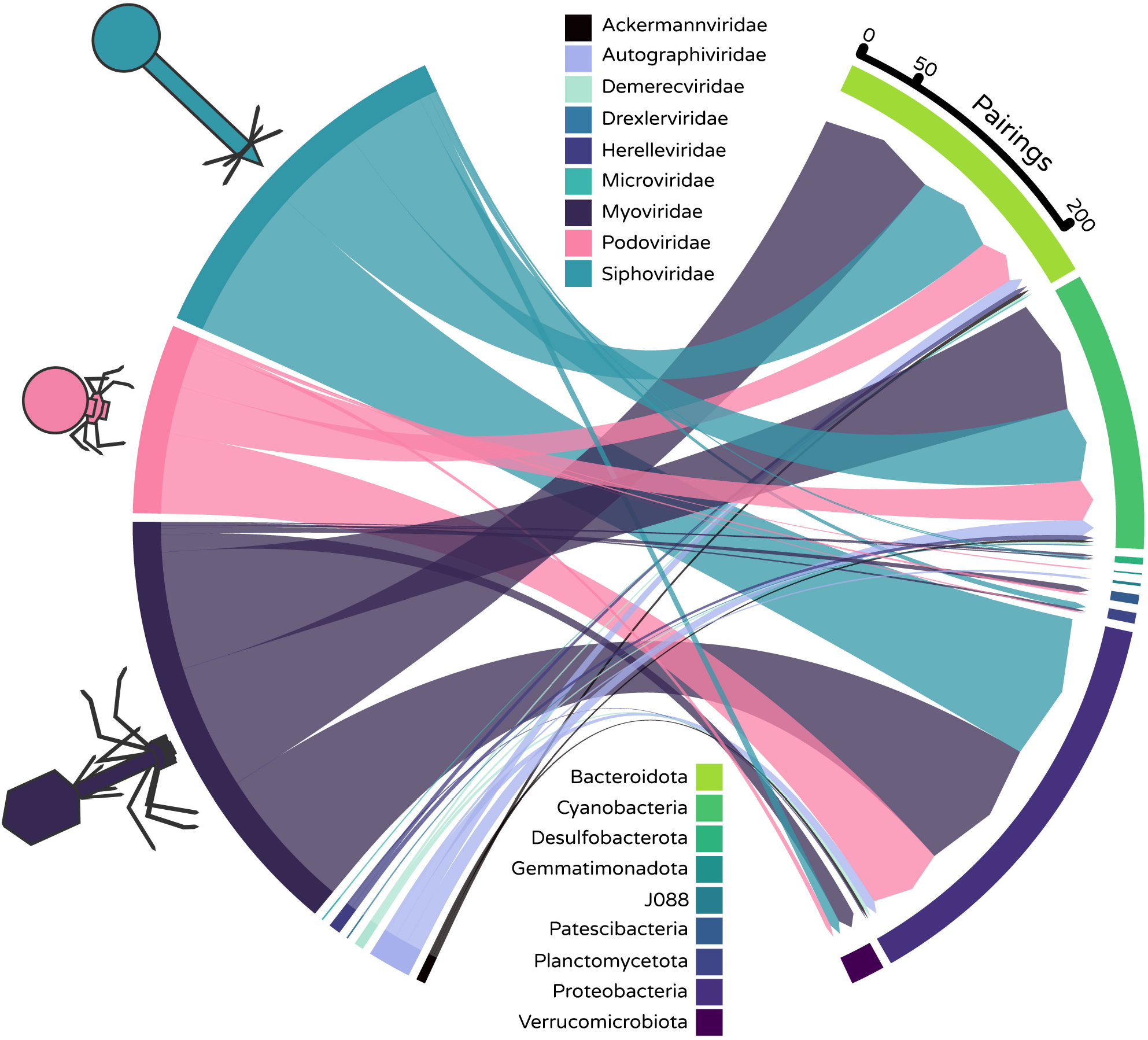
Diverse mat-associated viruses interact with diverse mat-building phyla. Interactions are illustrated using a directionally bipartite chord diagram of host-virus linkages with viral family (left) and host phyla (right). Bar size for each family or phylum encodes the total number of individual linkages (count) predicted from each viral family or for each host phylum.

### Cyanobacterial mats are characterized by an increasingly top-heavy trophic structure

To better understand patterns in the dynamics of predicted lineage-resolved host:viral pairings, patterns in the abundances of predicted hosts and their viruses in metagenomes were explored. Patterns were explored at the level of host phylum and host class. Phyla-resolved results are presented in main text figures (Figs. 4 & 5) – class-resolved results are presented in the appendix (Figs. S2 & S3). Ratios among viral and host coverages (VMR) in metagenomic read space consistently exceeded a 1:1 ratio and differed significantly by host phyla (Type II χ^2^; chisq = 241.83, df = 14, p < 2e-16; Fig. 4a). This suggests many viruses recovered may be actively lysing their predicted hosts with differential top-down interaction strengths across mat building populations by broad host taxonomic affiliation. Among mat variation in VMR was minimal (random effect variance 0.006 ± 0.07SD), suggesting strong spatial (among mat community) conservation of these patterns in viral trophic linkage strengths. To explore host virus relationships in cyanobacterial mats further, the shape in the abundance pyramid across host abundances was resolved from power law scaling among log-log abundances of viruses and their hosts. The power law scaling coefficient indicates the shape of the curve on ordinary axes, describing how predator abundance changes with prey abundance (58). Log-log regression of linked virus-host abundances revealed an increasingly top-heavy trophic structure within benthic cyanobacterial mats as host abundance increases (WLS; k = 1.25, SE = 0.047, t = 26.5, p < 2e-16), with strong concordance in the abundance relationships among viruses and computationally inferred hosts (adjusted r^2^ = 0.865). This further suggests bacteria and archaea experience significant top-down control from viral predation in cyanobacterial mats.

**Figure 4.**
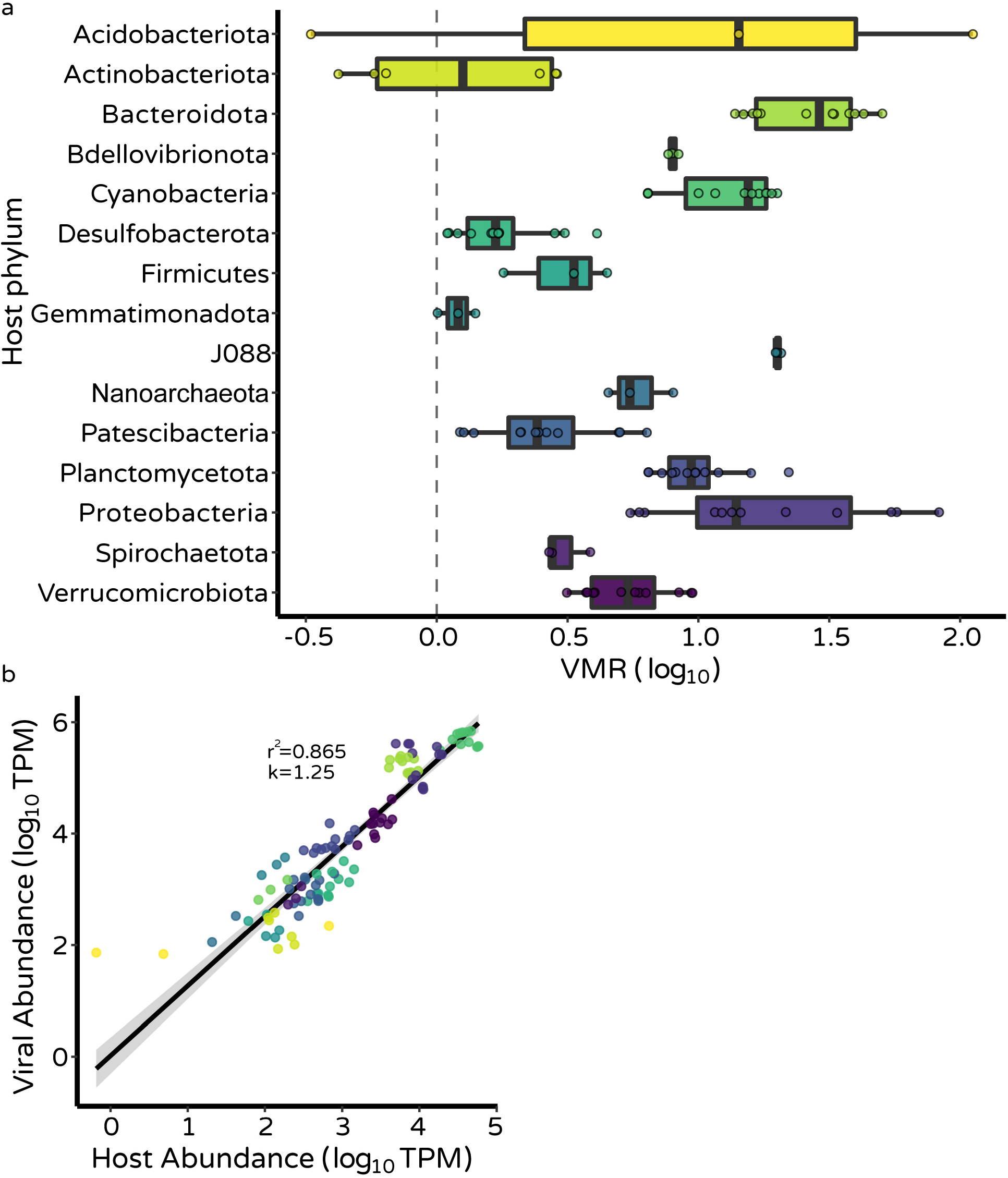
Cyanobacterial mats are characterized by a top-heavy trophic structure. a, Boxplot of VMR (log10 transformed) grouped at the level of host phylum showing ratios in metagenomic coverage consistently exceeding 1:1 (0 on log scale; vertical dashed line) b, Log-Log regression of viral and host abundances demonstrating an increasingly top heavy trophic structure as host abundance increases.

These patterns were well recapitulated when interrogating host class-level lineage-resolved linkages (Fig. S2). Among host class differences in VMR were detected (Type II χ^2^; chisq = 305.35, df = 21, p < 2e-16; Fig. S2a) similar to among phyla differences discussed above. Log-log power law scaling similarly predicted an overall increasingly top heavy trophic structure with respect to virus:host trophic linkages (WLS; k = 1.18, SE = 0.045, t = 26.4, p < 2e-16; Fig. S2b), with an adjusted r^2^ of 0.82.

### Metatranscriptomic recruitment suggests active predation across diverse hosts

To complement patterns observed among abundances of linked viral:host pairs in metagenomic read space, we developed and explored a similar metric in metatranscriptomic read space to traditional VMR which we term VMAR. VMAR leverages the changes in host transcriptional regulation associated with active viral infection and replication (4, 59), while directly assessing expressed viral activity (i.e. inherently subsets data and ecological interpretation to active infections). To contextualize patterns in VMAR with patterns from traditional VMR, we fit a WLS regression to logVMAR against logVMR. Patterns in VMAR and VMR were strongly concordant, suggesting that VMAR well captures dynamics inferred from more traditional measures alongside increased resolution of discerning active vs. inactive infections (adjusted r^2^ = 0.48; k = 0.84, SE = 0.08, t = 10.1, p < 2e-16; Fig. 5b). Viruses infecting all host phyla recruited reads from metatranscriptomes, indicating that hosts from the full diversity of mat-building phyla experience active viral infection. Mean VMAR differed significantly across host phyla (Type II χ^2^; chisq = 992.5, df = 14, p < 2e-16; Fig. 5a), suggesting differences in the interaction strength of actively infecting viruses by host taxonomy. VMAR estimates were unsurprisingly (because this represents the transcriptionally active subset) consistently higher than VMR estimates (Fig. 4 & 5), reinforcing that the cyanobacterial mat virome actively infects predicted bacterial and archaeal hosts. Power law scaling among log-log viral and host expression were explored as in metagenomic abundance described above, and revealed an even faster accelerating top-heavy relationship among expressed viral and host activity at increasing levels of host expression (OLS; k = 1.51, SE = 0.05, t = 30.6, p < 2e-16; Fig. 5c). This model similarly predicted strong concordance in patterns among viral and host expression (adjusted r^2^ = 0.8352; Fig. 5c).

**Figure 5.**
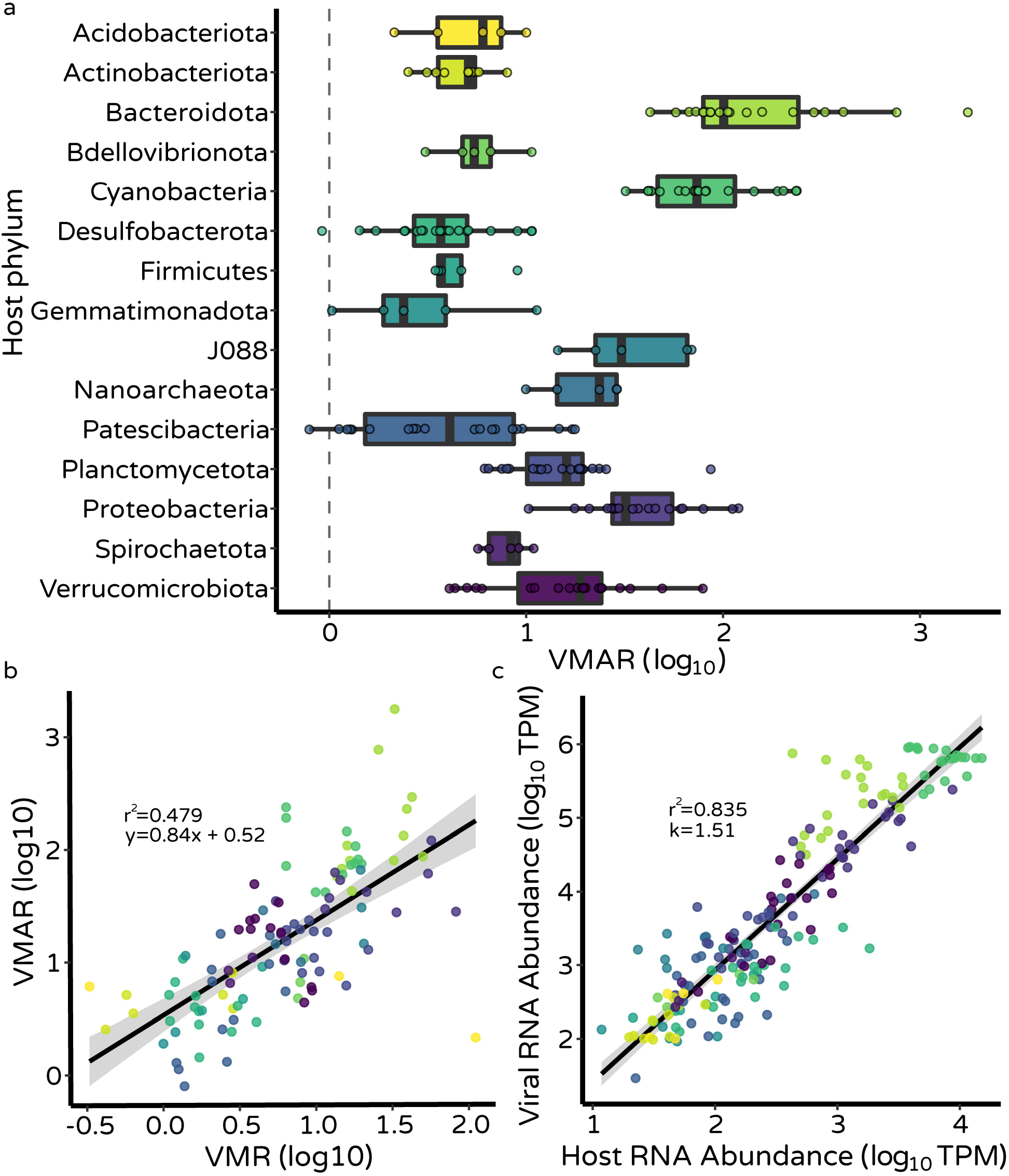
Cyanobacterial mats are characterized by a top-heavy trophic structure in transcriptionally active viruses. a, Boxplot of VMAR (log10 transformed) grouped at the level of host phylum showing ratios in metagenomic coverage consistently exceeding 1:1 (0 on log scale; vertical dashed line) b, Regression of VMAR against VMR showing strong concordance in ratios of virus:host activity and abundance. c, Log-Log regression of viral and host activity demonstrating an increasingly top heavy trophic structure as host abundance increases.

Patterns in viral activity were further explored at the level of host class, with significant differences in VMAR among host classes (Type II χ^2^; chisq = 890.9, df = 21, p < 2e-16; Fig. S3a). Power law scaling among log-log viral and host (class level) expression were explored as described above, and offer further evidence of an increasingly top-heavy trophic structure within the active viral subset of the cyanobacterial mat virome (OLS; k = 1.42, SE = 0.05, t = 30.0, p < 2e-16; adj. r^2^ = 0.7762; Fig. S3b).

Interestingly, global VMAR was temporally consistent across the sampled diel cycle (Type II χ^2^; chisq = 4.79, df = 4, p = 0.31), suggesting a general temporal consistency in active top-down interactions between viruses and microbial predators. Because there was not enough statistical power to test for the full interaction among host taxonomic affiliation and sampling time, data were subset to only include Cyanobacteria and putative infecting cyanophages. VMAR among subset cyanophages and host Cyanobacteria significantly differed across the sampled diel cycle (Type II χ^2^; chisq = 18.896, df = 4, p = 0.0008; Fig. S4).

## DISCUSSION

Here, we deeply sequenced longitudinally-collected paired metagenomes and metatranscriptomes from across a complete diel time-series to explore the diversity and structure of the coral reef benthic cyanobacterial mat virome. Using these sequencing data and leveraging previously curated MAGs assembled from this dataset, we further sought to interrogate the ecology of lineage-resolved host:virus interactions, with a specific focus on better understanding predation pressure from viruses in cyanobacterial mats. This sequencing effort represents one of the first in-depth explorations of viral ecology in a coral reef benthic cyanobacterial mat (38, 50), substantially expanding our knowledge on the genetic diversity and ecology of benthic marine environment viromes while contributing a refined public database of curated vOTUs consisting of recovered dsDNA, ssDNA, and dsRNA viruses (the vMAT database) from a critically unexplored benthic habitat for further exploration and comparison with other environmental datasets obtained in future sequencing endeavors from other diverse environments. Overall, our data suggest that viruses are highly active members of the cyanobacterial mat community, and likely exert strong top-down control on bacterial and archaeal population abundances.

Our finding of such strong evidence for relatively systemic top-down control across host taxa in cyanobacterial mats is surprising (Fig. 4, 5). Previous assessments of fast-growing, high-density microbial systems have suggested an overwhelming predominance of lysogenic interactions over lytic interactions (9, 26, 60), with few exceptions (61–63), which support our initial hypothesis of a weak top-down signal in similarly fast-growing high-density cyanobacterial mat communities. Indeed, aquatic benthic systems in general have long been thought to have generally low rates of bacterial mortality from infecting viruses, despite high mortality in surrounding water column communities (64, 65). While we did not explicitly interrogate ratios of lysogeny vs. lysis in this system, the VMRs and VMARs documented herein would be highly unlikely in a system dominated by lysogeny, as we would expect VMR and (to a lesser extent) VMAR to be closer to 1:1 (60). Additionally, the shape of the relationships between log-log viral host abundance/expression were both upward curved on traditional axes, pointing to increasing viral control as host abundance and activity increase (Fig. 4, 5). Many phyla-specific and class-specific prey abundance gradients span ∼1 or less order of magnitude, which unfortunately precluded us from meaningfully resolving phyla-specific host-viral power law behavior, or from effectively interrogating the relationship between host abundance and VM{A}R (i.e. [x/y]/y), which would be useful for further analyzing the relationship between viral ecology and host abundance in this system. Taken together, these results highlight 1) the dire need to incorporate patterns from diverse contemporaneously unexplored microbial systems into theory and quantitative frameworks linking describing viral influence on microbial communities, and 2) the need to understand how biotic and abiotic context can drive fine-scale nuance in virus-microbe trophic interactions and contribute to patterns that depart from, and overwhelmingly subvert, general predictions from theory (53).

Viral sequences recovered in this study were highly divergent from reference viruses, and previously curated viruses from seawater and permafrost (Fig. 1, 2). Strong endemism of benthic viruses has previously been demonstrated in deep sea sediment viral communities (63), and suggest that marine benthic viromes may generally harbor strongly unique viral communities from previously explored environments. Many of the viral sequences reconstructed herein likely belong to novel taxonomic groups with no current representatives in viral databases, as a significant proportion of vOTUs could not be assigned to a taxonomic family (Fig. 1). Large gaps in unexplored diversity present in this benthic viral dark matter remain and warrant detailed exploration.

Strong lineage-specific patterns in intra-mat trophic structure (Figs. 4, 5, S2, & S3; potentially coupled to host trophic affiliation [53]) may in-part contribute to patterns of spatial genomic diversification among mat-building populations (Cissell & McCoy in review), and likely impact intra-mat chemistry and nutrient recycling (66). Bacteria-phage coevolution is known to increase both phage and bacterial divergence rates mediated via reciprocal adaptation (evolutionary conflict), and promote diversity across multiple levels of ecological organization, affecting overall community structure (12). In this way, strong viral activity likely directly (via biomass turnover and cell lysis) and indirectly (via metabolic augmentation and expansion of physiological potential [virocells]) helps drive cyanobacterial mat nutrient cycling. Temporal niche partitioning and metabolic specialization among host taxa (Cissell & Mccoy in review) could interact with observed viral activity via physiology and growth-phase mediated effects (67), which may cause differential shifts in stoichiometry across a diel cycle given the dramatically different active metabolic processes at day vs. night (Cissell & McCoy in review). While there was limited evidence for a diel signal in overall viral activity across sampling time points (Fig. S4), it may be that diel variability in infectivity was obfuscated by this sweeping aggregation across temporally specialized communities. Characterizing cyanophage:Cyanobacteria VMAR specifically revealed reduced VMAR when cyanobacterial growth likely peaked (Fig. S4; Cissell & McCoy in review). However, graphical exploration of temporal patterns across host phyla suggests that the strongest temporal signal in VMAR is within the phyla Cyanobacteria, with relative temporal consistency across other phyla (Fig S5). This is supported by the lack of a significant trend in virome Bray-Curtis dissimilarity along a gradient of increasing temporal distance (GAM, edf = 1.3, F = 0.15, p = 0.8, Deviance = 2.05%; Fig. S6), and no clear trend in evenness of viral assemblage activity toward dominance across time (Fig. S7). Cyanobacteria are well known for exhibiting strong biological rhythmicity across light-dark cycles (response at the diel scale; 68), with similar dynamism in cyanophage infection dynamics mediated primarily via light-dependent adsorption rates, burst sizes, and host expression patterns (reviewed in 69). Better understanding fine-scale temporal variation in viral activity in response to temporal variability in host physiological state will be critical to scale emergent cyanobacterial mat community physiology to ecosystem scale chemistry, but will require datasets of substantial size to well-resolve an interaction term between host taxonomic identity and time.

These data highlight the ability of using host and viral sequences from the same dataset to recreate meaningful patterns in predator-prey ecology with viruses (17). This is especially useful in unexplored environments with few cultured representatives, such as those taxa in coral reef benthic cyanobacterial mats (38). Future laboratory experiments should leverage the computationally-predicted virus-host associations presented herein toward isolating and testing specific patterns in predator-prey or mutualistic interactions with the cyanobacterial mat virome in order to gain further functional insights into this diverse viral dark matter (70). Our data overall suggest that viruses exert considerable influence over the dynamics of mat building populations and likely are major players in modulating the functional ecology of cyanobacterial mats on reefs. Viruses should be increasingly considered when exploring patterns in the demography and ecophysiology of cyanobacterial mats toward holistically understanding the causes and consequences of mat proliferation on coral reefs (53).

## MATERIALS & METHODS

### Overview of sampling, sequencing, quality control, and metagenome assembly

Sampling strategy, library preparation, sequencing, data quality control, metagenome assembly, metagenome binning, and MAG taxonomic placement are described in detail elsewhere (Cissell & McCoy in review). For concise contextualization, relevant methods will be very briefly summarized here. Sampling was conducted in Bonaire, Caribbean Netherlands, at Angel City reef (12°06’12.2” N, 68°17’14.0” W) on SCUBA. Sampling was permitted under research permit RWS-2019/9554 issued by Rijkswaterstaat Ministerie van Infrastructuur en Waterstaat, and with special permissions from Stichting Nationale Parken (STINAPA) Bonaire. Four spatially distinct cyanobacterial mats at mean 16.6m depth were sampled (Table S1; 1.5mL volume each) at 5 evenly spaced time points (09:00, 15:00, 21:00. 03:00. 09:00) across a complete diel cycle beginning at 09:00 Atlantic Standard Time on 29/06/2019 and ending at 09:00 on 30/06/2019. A total of n=80 samples were collected – 4 samples per mat individual per sampling time – and stored in 2x their volume of DNA/RNA Shield (Zymo), frozen at -20°C while in the field (18 days), and at -80°C upon returning to Florida State University. Replicate samples from within a mat individual within a sampling time were pooled, had metagenomic (total DNA) and metatranscriptomic (cDNA from rRNA-depleted total RNA) libraries prepared, and were sequenced using paired-end 150bp chemistry (300 cycles) on an Illumina NovaSeq 6000 at Novogene, generating a total of ∼7.5 billion reads across all samples (Table S2, S3).

These sequencing reads are available in the NCBI sra database, accessible under BioProject accession number PRJNA632569.

Quality controlled metagenomic reads were coassembled within mat individual (n=3 read sets each) *de-novo* with *MEGAHIT* (Table S4; 68); binned unsupervised using *Vamb* ver. 3.0.2 (72), *Metabat2* ver. 2.15 (73), and *Maxbin2* ver. 2.2.7 (74); coalesced using *DasTool* ver. 1.1.2 (75); manually refined (supervised bin curation) using *Anvi’o* (76); and dereplicated at 99% identity using *dRep* ver. 3.2.0 (77), producing 261 medium-to-high quality MAGs. Taxonomy was assigned to MAGs using *GTDB-Tk* ver. 1.5.0 (78) against GTDB release 06-RS202, with select *BLASTn* phylogenetic taxonomic refinement.

### Metatranscriptome quality control and assembly

Raw cDNA reads were quality trimmed to a Phred quality score threshold of 20, minimum length threshold of 50bp, and had adapters removed using *TrimGalore* as previously described (Cissell & McCoy in review). *SortMeRNA* ver. 4.2.0 (79) was used to remove rRNA reads that were not removed during library preparation. Metatranscriptomes were subsequently coassembled *de-novo* from quality controlled paired-end metatranscriptomic read sets from the same mat individual (n=5 read sets coassembled per mat) using *rnaSPAdes* ver. 3.13.0 (80) using *k*-mer sizes of 49 and 73, retaining contigs of length ≥ 2.5kbp. Though quantitative assessment of ‘good quality’ is inherently difficult to define in complex mixed-assemblage reference-free *de-novo* metatranscriptomic assemblies, overall quality of metatranscriptome coassemblies was assessed using multiple quantitative assembly summary statistics generated using *METAQUAST* (Table S5; 78).

### Viral contig identification

Viruses (including phages integrated as prophages), were initially computationally defined in metagenomic (dsDNA and ssDNA viruses) and metatranscriptomic (RNA viruses) coassembly contigs based on recognized homologs to viral hallmark genes and enrichment of viral signature proteins (referenced-based definition) using *VirSorter2* ver. 2.2.3 (82), retaining predicted contigs ≥2.5kbp in length with a minimal score cutoff of 0.5. This more relaxed length threshold was imposed to increase classifier sensitivity within this highly novel virome environment. Prophage boundaries were defined and cleaned (host contamination removed, viral sequence length filter removed), and the quality (including completeness) of retained viral contigs was conservatively assessed using *CheckV* ver. 0.8.1 (83). A total of 20,351 contigs across all coassemblies were flagged as putative viral contigs. Open reading frames and Auxiliary Metabolic Genes (AMGs) were predicted and annotated on putative viral contigs using homology searches against PFAM, dbCAN, RefSeq viral, VOGDB and MEROPS implemented in *DRAMv* ver. 1.2.2 (84) using default parameters. Viral contigs were subsequently conservatively manually curated using custom scripts and supervised screening based on combinations of viral and host gene counts from *CheckV*, hallmark gene counts from *VirSorter2*, contig score, and gene-level annotations from *DRAMv* following previously benchmarked protocols (85) to remove potential false positive predictions (broadly screening for enrichment in viral-like or viral hallmark genes; depletion in host genes), retaining 12,843 total contigs from metagenomes, and 214 total contigs from metatranscriptomes. Manually curated dsDNA, ssDNA, and RNA viral sequences were clustered at 95% minimum sequence identity within coassembly using *CD-HIT-EST* ver. 4.7 (86), creating non-redundant_95%_ databases with vOTUs that represent a combination of free viruses, integrated prophages, and actively infecting viruses (vMAT database). Clustered vOTUs were further validated using non-reference-based HMM similarity searches implemented in *VIBRANT* ver. 1.2.0 (87) using the -virome flag, with a mean 74% ± 1.6% SD and 21% ± 9.0% SD of all DNA and RNA curated viral contigs validated, respectively (Dataset S1). It should be noted that *VIBRANT* performs relatively poorly (sensitivity) on RNA phages in comparison to *VirSorter2*, and so this low validation on predicted RNA viral sequences is not surprising.

### Integrative viral taxonomic assessment and environmental similarity network reconstruction

Broad assemblage-scale taxonomic context (family) was assigned to viral contigs by integrating two independent machine learning-based methods. First, family-level taxonomy was assigned with translated sequence similarity searches utilizing reciprocal *DIAMOND BLASTp* mapping and protein Markov clustering against the Caudovirales Proteins database, followed by subsequent training of convolutional neural networks implemented in *PhaGCN* ver. (88).

Genomic protein-sharing networks were also created among all nonredundant viral contigs reconstructed in this study and all sequences in RefSeq_88_ by first creating amino acid translations of ORFs on phage nucleotide sequences using *Prodigal* ver. 2.6.3 (89) followed by inferring viral cluster network structure using *vConTACT2* ver. 0.9.19 (90) with *DIAMOND BLASTp* and default parameters for further resolving family-level taxonomy and for visualizing viral cluster network topography. Betweenness centrality topology of resulting network edges was calculated as sum(g_iej/g_ij, i != j). For resolving taxonomy of those vOTUs sharing significant protein homologs with clusters that are composed of multiple representative families in RefSeq (n = 2 screened clusters), edge weight was used to manually resolve best supported taxonomy. Among viral clusters, *vConTACT2* and *PhaGCN* had concordant taxonomic predictions on n = 24 vOTUs, and discordant taxonomic predictions for n = 3 vOTUs. In the latter cases, *PhaGCN* taxonomy was retained. To leverage the independent strengths of each prognostic method against the shared diversity among mat samples, vOTUs that did not cluster with known RefSeq viral genomes and that could not be assigned taxonomy in *PhaGCN*, but formed clusters with at least one other cyanobacterial mat vOTU that had been assigned taxonomy by *PhaGCN* with a confidence of ≥ 0.75 (mean confidence score 0.97 ± 0.06SD) were assigned the corresponding representative family-level taxonomy (n = 339). Many of these clusters possessed numerous independently concordant *PhaGCN* taxonomic predictions, increasing our confidence in assigning all vOTUs within these clusters to a conservative shared family-level taxonomy (viral clusters can be thought of as approximating genus-level relatedness (90)).

To contextualize and better understand how the benthic viral contigs generated in this study compare to other relevant environmental meta’omics-derived viral contigs, an additional genomic protein-sharing network among the viral contigs from this study (vMAT), viral contigs reconstructed from permafrost (n=1,907 populations; GenBank accession: GCA_003191745.1; (17)), and all of the GOV 2.0 (23, 28, 91) epipelagic and mesopelagic seawater viral populations (retrieved from iVirus; n=195,728 contigs) was created using *vConTACT2* with default parameters. Translated ORFs were obtained with *Prodigal* as described above. Betweenness centrality topology of resulting network nodes was calculated as above for edges.

### Integrative host prediction

Putative virus-host interactions were predicted by integrating predictions from multiple computational host prediction strategies using the refined viral contig database, including the following approaches: CRISPR-spacer::protospacer matching, host-viral homology matching, and oligonucleotide frequency screening. Because the custom host query database used consists only of prokaryotic MAGs (Bacteria and Archaea), downstream infection dynamic analyses necessarily focus exclusively on the phages and Archaeal viruses found in the viral contig database and exclude any viruses infecting the limited (from relative abundance) Eukaryotic members of the mat community (38; Cissell & McCoy in review). CRISPR spacers were predicted using *MinCED* ver. 0.4.2 (92) on a custom database of nonredundant (99%) manually refined MAGs reconstructed from benthic cyanobacterial mats (Cissell & McCoy in review).

Identified CRISPR spacers were packaged into a custom nucleotide database against which viral contig protospacers were subsequently queried using *BLASTn* using previously benchmarked optimum search standards for virus-host prediction across the length of the spacer (*BLASTn* - short; maximum E-value 1; gap opening penalty 10; gap extension penalty 2; mismatch penalty 1; word size 7; Ref 79), conservatively retaining best hit pairs with a stringent allowance of only ≤1 mismatch (94). Additionally, viral contigs were compared to refined MAGs using *BLASTn* with the following criteria: ≥70% minimum nucleotide identity, ≥75% coverage over length of viral contig, ≥50 bit score, and ≤0.001 e-value (63). Finally, shared *k*-mer frequencies (25-mers) among viral contigs and refined MAGs were compared using *PHIST* ver. 1.0.0 (95), retaining predicted pairs with adjusted p-values <0.001 and a greater number of *k*-mer matches than the study-wide median (36 matches). The single best hit was retained from each method; to resolve multiple host predictions, virus-host linkages supported by multiple approaches were retained. If no consensus among all approaches was identified, the following ranked criteria were used (63): (i) best CRISPR spacer match; (ii) nucleotide sequence homology; (iii) oligonucleotide frequency comparison. The quality of virus-host pairings were subsequently further screened via manually comparing viral gene annotations with predicted hosts (ex: phages with PSII protein D1 [*psbA*] and/or protein D2 [*psbD*] or enrichment in characterized cyanophage protein hits predicted to have cyanobacterial host).

### Viral abundance and activity quantification

Copies Per Million (CPM) values (96) were calculated using custom scripts pulling raw counts and lengths using CoverM from recruitment profiles generated with *Bowtie2* ver. 2.3.5.1 (97), and were used as proxies for relative abundances of viruses and their hosts for calculating Virus-Microbe Ratios (VMRs; Total Phage + Host add to 1e6). Viral abundance profiles were only calculated on DNA viruses. Similarly, RNA reads were globally queried against host genomes and vOTUs to generate activity profiles using *BWA-MEM* ver. 0.7.17 (98). Virus-Microbe Activity Ratios (VMARs; Total Phage + Host add to 1e6) were subsequently generated from Transcripts Per Million (TPM) normalized counts pulling high-quality RNA alignments to annotated gene-level features. VMAR leverages the cytopathogenesis of viral infection, namely global inhibition of host expression (4, 59), for ecological interpretability. TPM was calculated from summed raw feature counts per genome using the summed length of predicted ORFs (*Prodigal*) for length normalization rather than total genomic length. Activity profiles were not generated for RNA viruses.

### Statistical procedures

All statistical analyses were conducted using R ver. 3.6.2 within RStudio ver. 1.2.5033. Data visualizations were created using either R::ggplot2(ver. 3.3.3), R::ggraph(ver. 2.0.5), or R::circlize(ver. 0.4.13). Model assumptions for all models, unless otherwise specified, were assessed from model residuals graphically using R::DHARMa. To quantify a community-wide power law scaling parameter between viral and host relative abundances, we fit Weighted Least Squares (WLS) linear regressions to log10 transformed viral and host relative abundances normalized to CPM, where the most appropriate weighting structure (W) was derived from graphical exploration of residual patterns and given as the inverse of W=|ε|∼y^2^ using residual structures extracted from simple Ordinary Least Squares (OLS) regressions fit with the same predictor and response variables. The assumption of homoskedasticity was still violated following weighted regression, however WLS coefficient estimates remain robust against heteroskedasticity (unbiased estimators). Error structures presented herein are from the fit WLS models and should be interpreted carefully. To better understand how VMR varies by host taxonomic affiliation, we fit random-intercept linear mixed effects models (lme4::lmer) using Restricted Maximum Likelihood (REML) to log_10_ transformed VMR (log_10_[P/H]) against a fixed effect of either phyla or class (below class taxonomy was generally not informative in these MAGs with few exceptions) where the intercept varies by spatially distinct mat. Log_10_ transformed relative abundances (CPM) were preferred in these analyses to centered log ratio or isometric log ratio transformed counts to facilitate easier data integration and comparison with preexisting literature values (9, 60, 99).

To complement patterns characterized using DNA-based evidence, expressed viral activity was assessed from RNA recruitment against gene-level features. To explore the relationship between VMAR and VMR, a WLS model was fit to log_10_ transformed VMAR (from TPM normalized viral / host expression relationships) against log_10_ transformed VMR (calculated as above) with weighting structure defined as above to extract goodness of fit (r^2^) and significance of relationship (from p). Spatial variability was packaged into the random effects structure of an additional random-slopes linear mixed effects model fit using REML with an identical fixed effects structure to the OLS model while allowing for the intercept and slope to vary across spatially distinct mats with respect to logVMR. There was strong concordance in fixed effect coefficient estimations between the OLS and mixed effects model. An OLS regression was fit to log_10_ transformed viral and host RNA abundances normalized to TPM to explore power-law scaling among viral and host activity. To better understand temporal and host phyla-specific patterns in VMAR, a linear mixed effects model was fit to log_10_ transformed VMAR with both sampling time point and host phyla fit as fixed predictors where both the slope and intercept were allowed to vary by sampled mat with respect to host phylum. Further, activity data were subset to only include putative Cyanophage-Cyanobacteria interactions, and temporal patterns from this subset were interrogated in a linear mixed effects model fit to log_10_ transformed VMAR with sampling time point as the fixed predictor and random intercepts fit to sampled mat. Model assumptions were assessed graphically as above.

## ACKNOWLEDGMENTS

We thank R.L. Francisca and C.E. Eckrich at Stichting Nationale Parken (STINAPA) Bonaire and Rijkswaterstaat Ministerie van Infrastructuur en Waterstaat for research permissions for conducting this work within the Bonaire National Marine Park. We extend our thanks to J.C. Manning, I. Basden, M. Dziewit, and B. Clark for assistance in the field. We thank A. Brown and the Florida State University (FSU) Department of Biological Science Core Facility for help with library preparation. We thank C. Peters for supporting diving logistics at FSU. We thank D.K. Okamoto for help with statistical analyses. Finally, we thank J.R. Cissell, K.M. Majzner, J.C. Manning, K.L. Dobson M. Huettel, D.K. Okamoto, K.M. Jones, and D.R. Rokyta for discussions and feedback on earlier versions of this manuscript.

Funding for this work was provided by the the Phycological Society of America (Grant-In-Aid of Research), PADI Foundation (Grant Number: 40829), and the Western Society of Naturalists (Rafe Sagarin Fund for Innovative Ecology) awarded to E.C. Cissell. Additional funding was provided by a gift from the Tatelbaum Fund and start-up funding from FSU to S.J. McCoy. E.C. Cissell was supported by a National Science Foundation Graduate Research Fellowship (Grant Number: 074012-520-044116) and a Mote Research Assistantship from the William R. and Lenore Mote Eminent Scholar in Marine Biology Endowment at FSU.

## Notes

### Competing Interest Statement

The authors have declared no competing interest.

