## Appendix S1 for "Top-heavy trophic structure within benthic viral dark matter"

### Supporting Information for
### Top-heavy trophic structure within benthic viral dark matter

Ethan C. Cissell[1*] & Sophie J. McCoy[1]
[1]Department of Biology, University of North Carolina at Chapel Hill, Chapel Hill, NC, USA

*Address correspondence to: Ethan C. Cissell

**This PDF file includes:**

Figures S1 to S7
Tables S1 to S5

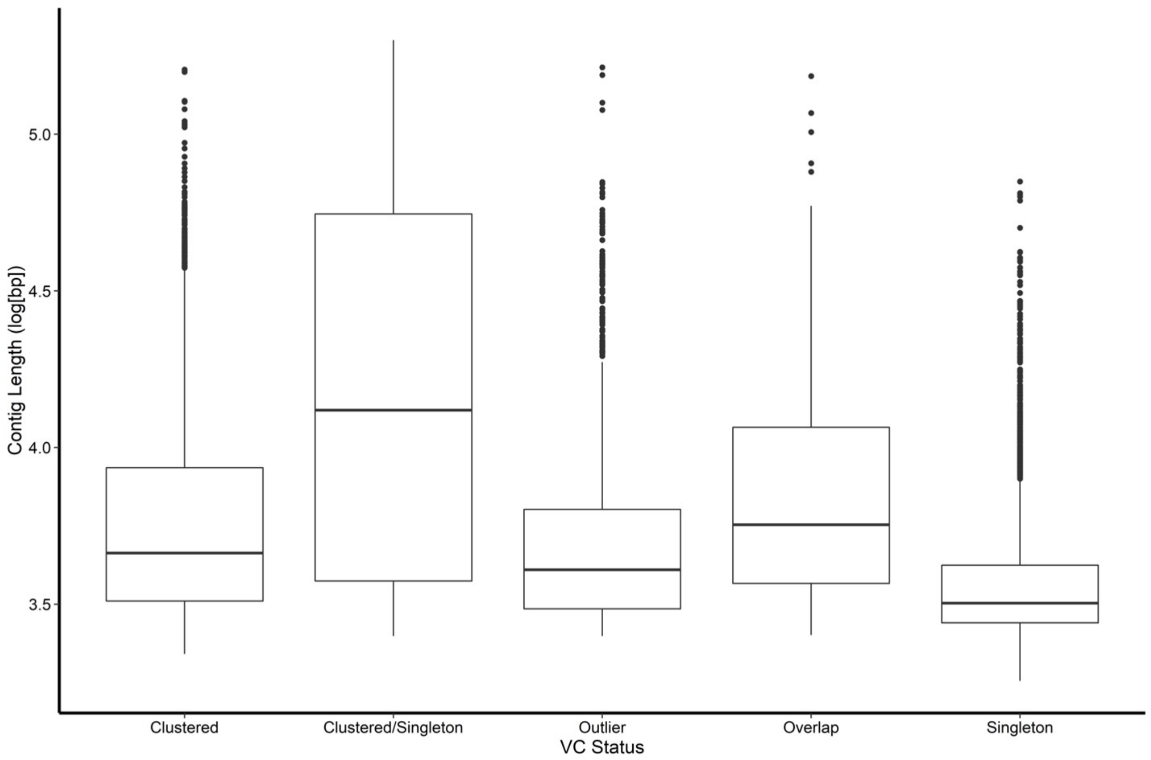


**Fig. S1.** Boxplots (median and interquartile range) of the relationship between viral genome length ($log_{10}$bp length) and clustering status. Individual points denote outliers.

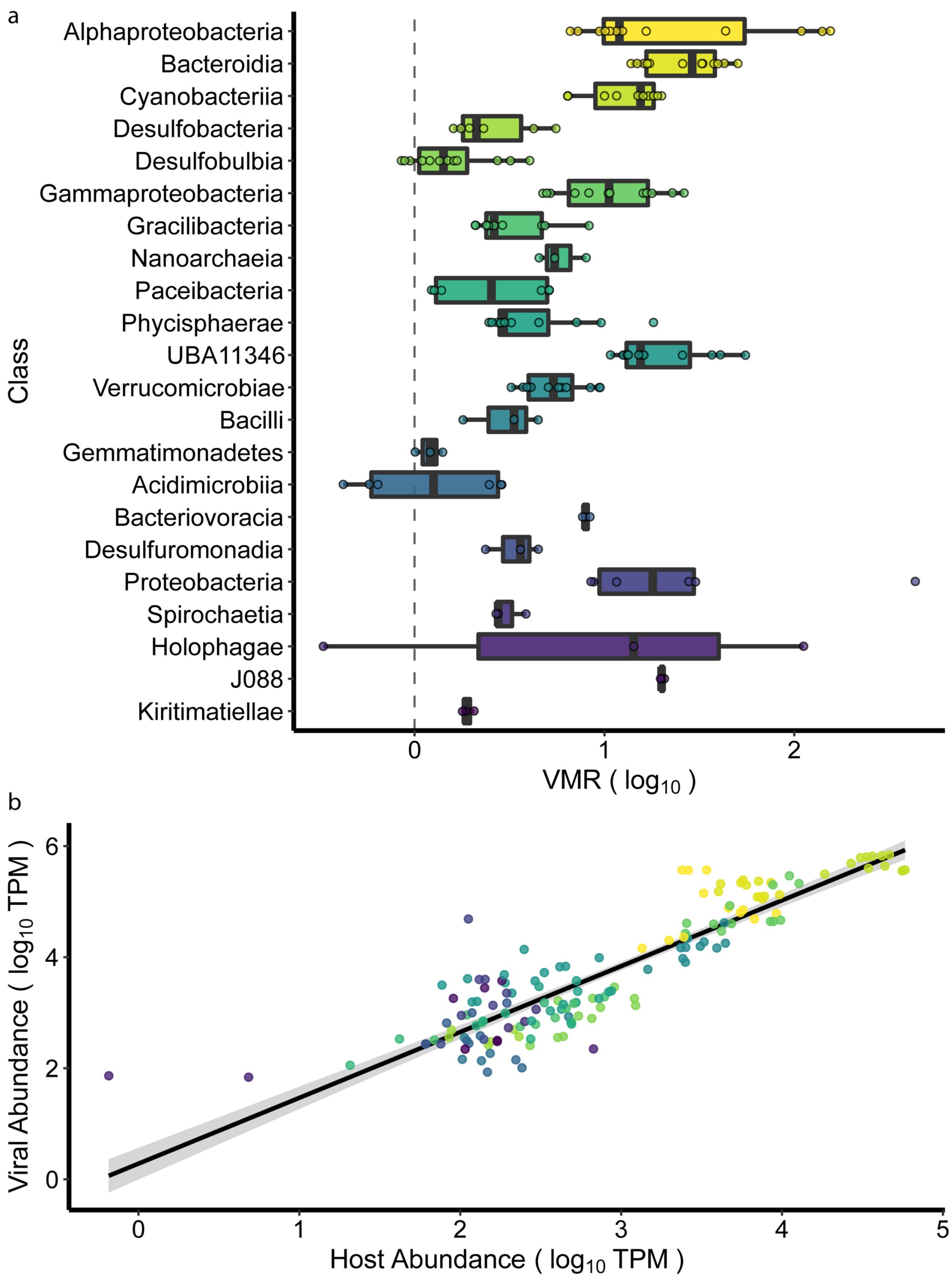


**Fig. S2.** Cyanobacterial mats are characterized by a top-heavy trophic structure. a, Boxplot of VMR (log10 transformed) grouped at the level of host class showing ratios in metagenomic coverage consistently exceeding 1:1 (0 on log scale; vertical dashed line) b, Log-Log regression of viral and host abundances demonstrating an increasingly top heavy trophic structure as host abundance (class summarized) increases (WLS; k = 1.18, SE = 0.045, t = 26.4, p < 2e-16 $r^2$ = 0.82).

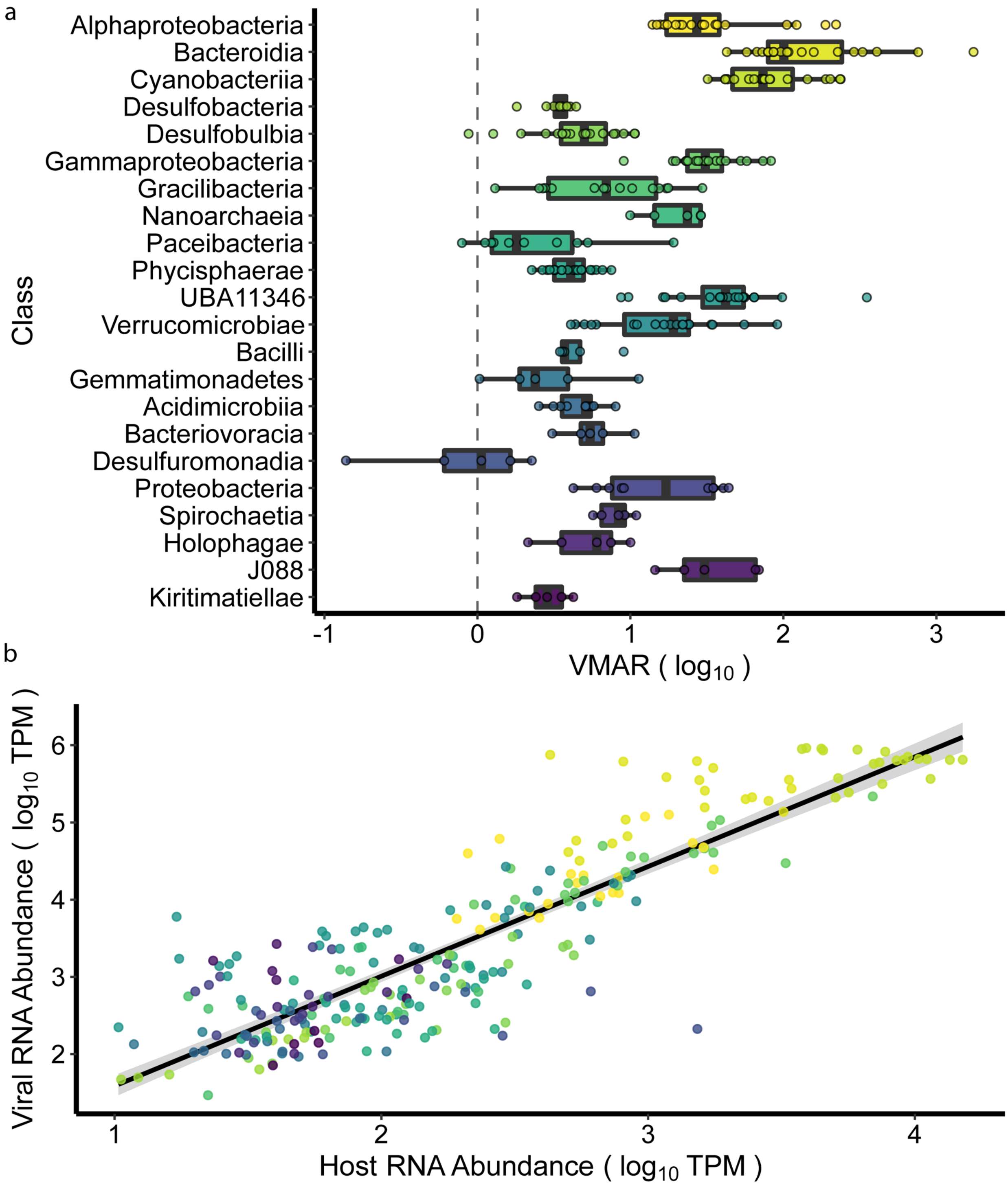


**Fig. S3.** Cyanobacterial mats are characterized by a top-heavy trophic structure in transcriptionally active viruses. a, Boxplot of VMAR (log10 transformed) grouped at the level of host class showing ratios in metagenomic coverage consistently exceeding 1:1 (0 on log scale; vertical dashed line) b, Log-Log regression of viral and host activity demonstrating an increasingly top heavy trophic structure as host abundance increases (OLS; $k = 1.42$, $SE = 0.05$, $t = 30.0$, $p < 2e-16$; adj. $r^2 = 0.7762$).

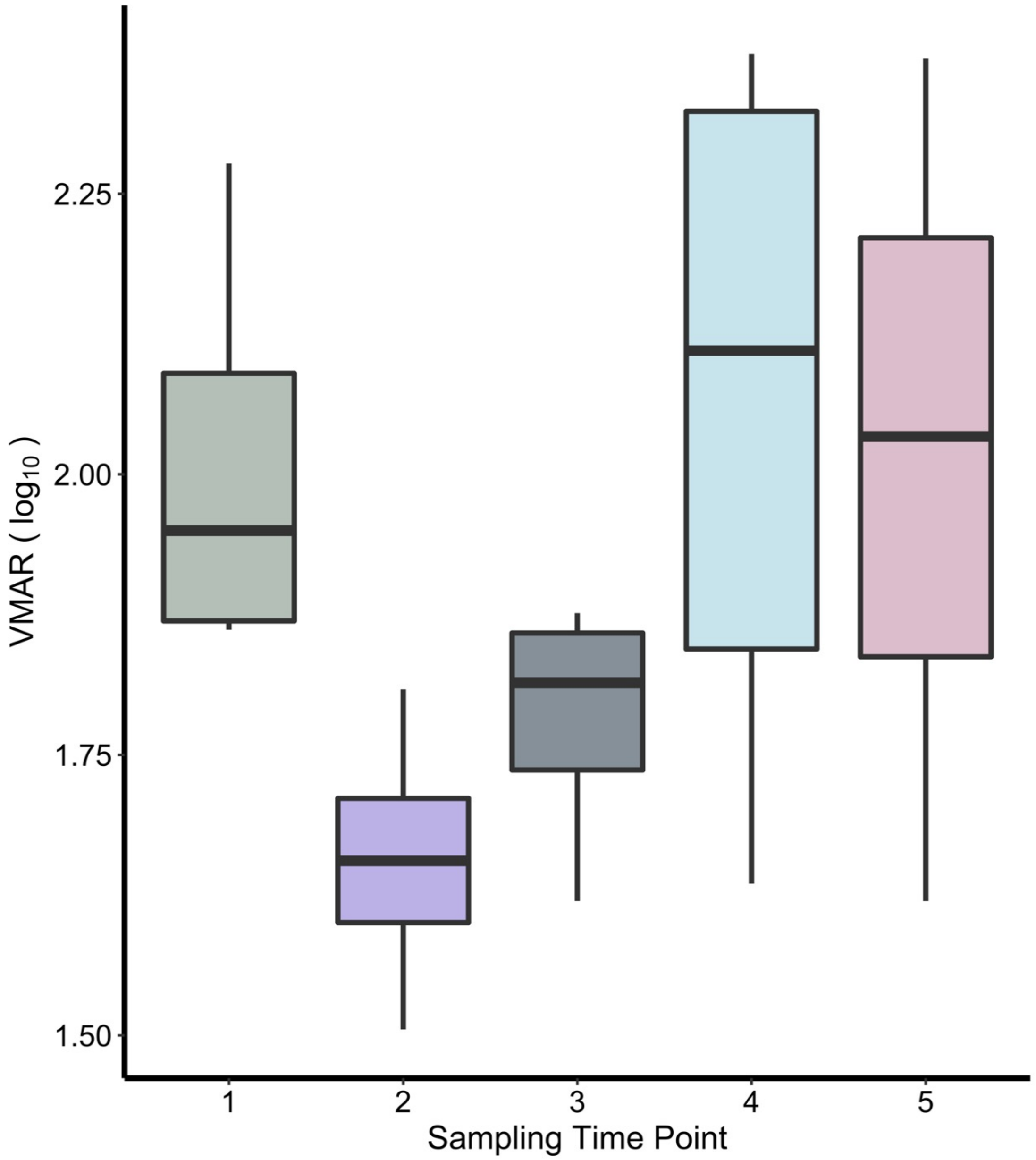


**Fig. S4.** Boxplots of VMAR ($log_{10}$ transformed) among putative cyanophage and Cyanobacteria grouped by sampling time point across the diel cycle. VMAR of cyanophage:Cyanobacteria pairs displayed a significant diel signal (Type II $\chi^2$; chisq = 18.896, df = 4, p = 0.0008).

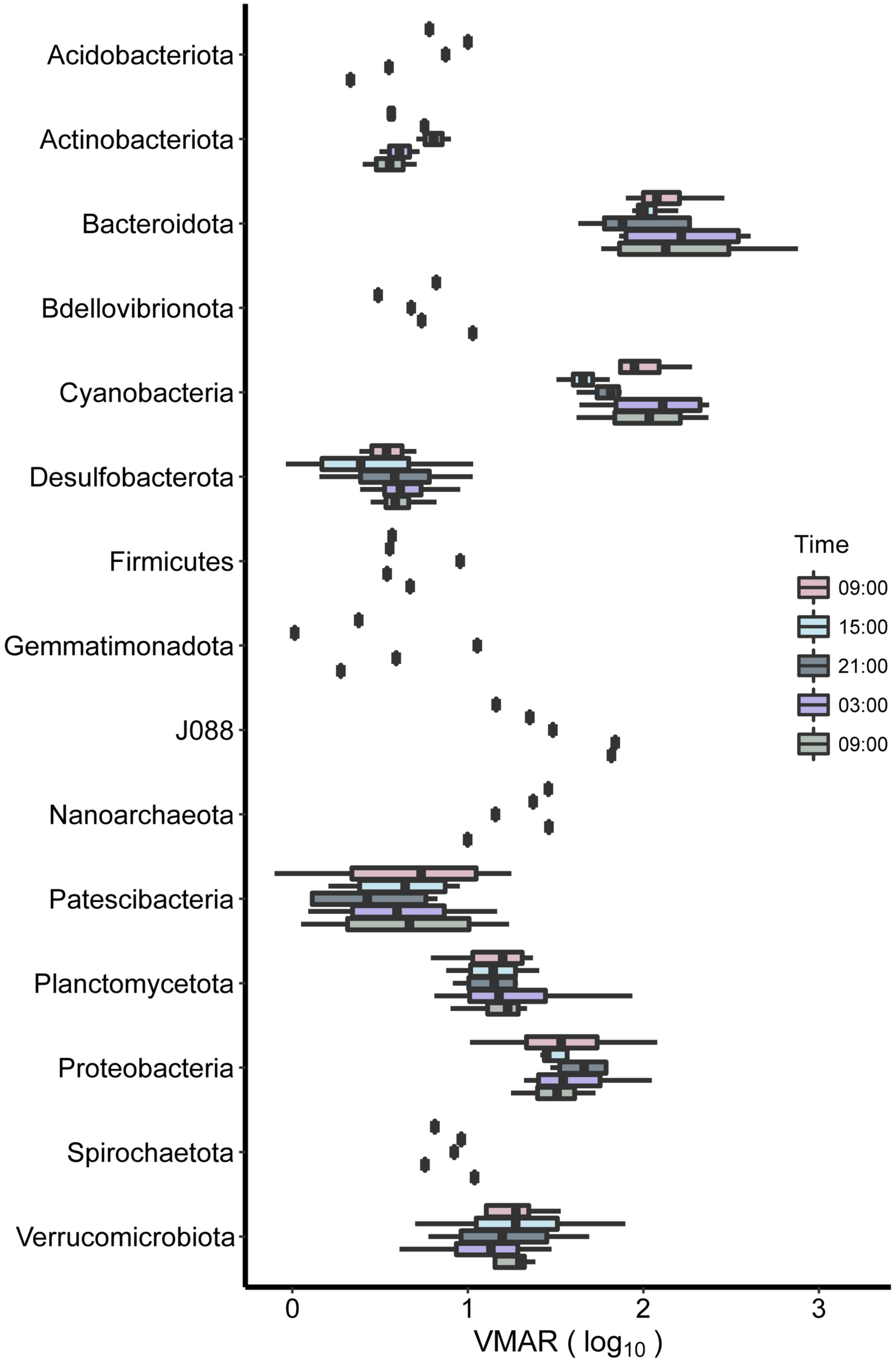


**Fig. S5.** Boxplots of VMAR ($log_{10}$ transformed) among virus:host pairs grouped at the level of host phyla and by sampling time point across the diel cycle.

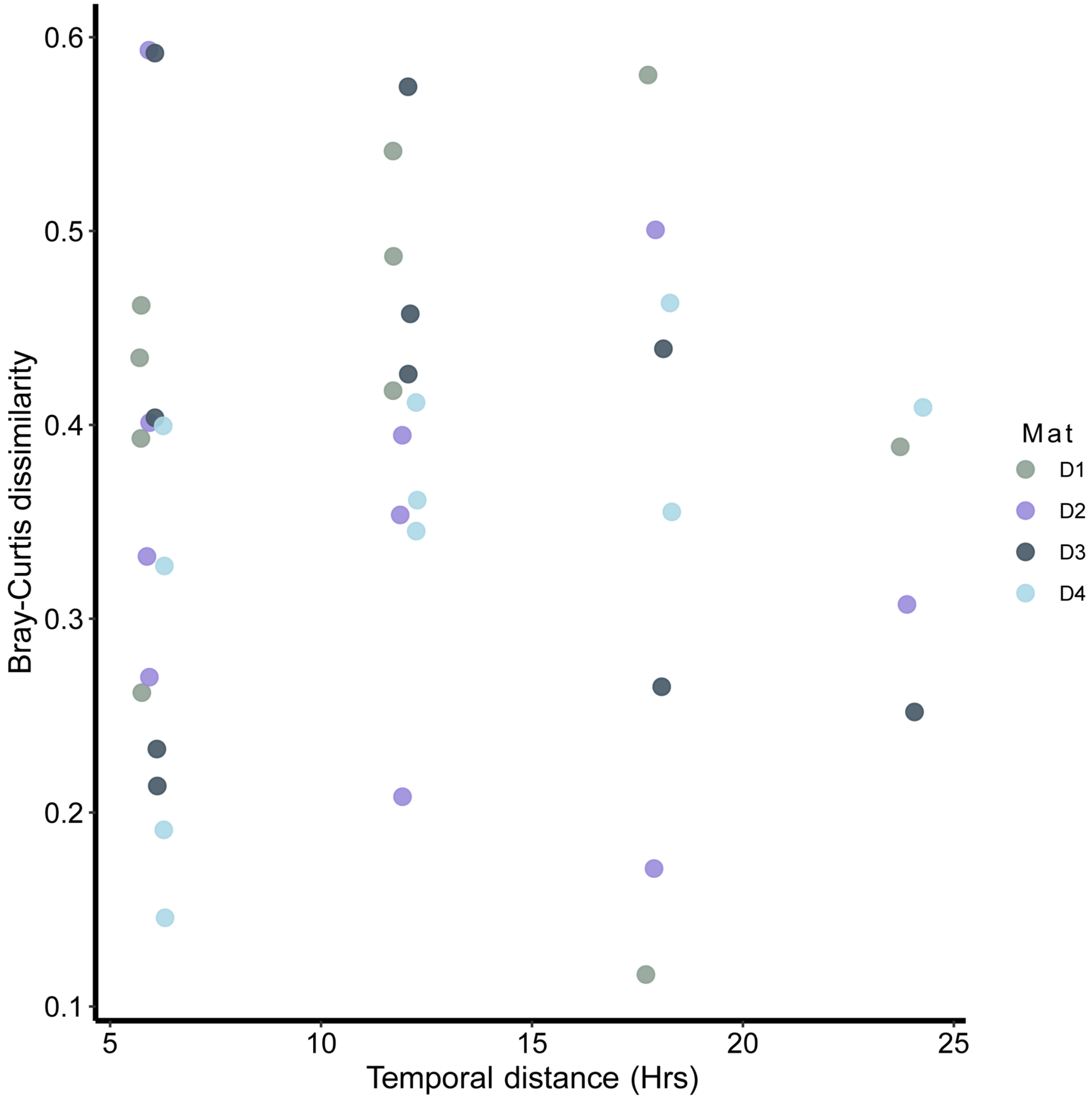


**Fig. S6.** Scatter plot of Bray-Curtis dissimilarity across temporal distance between samples (in hours).

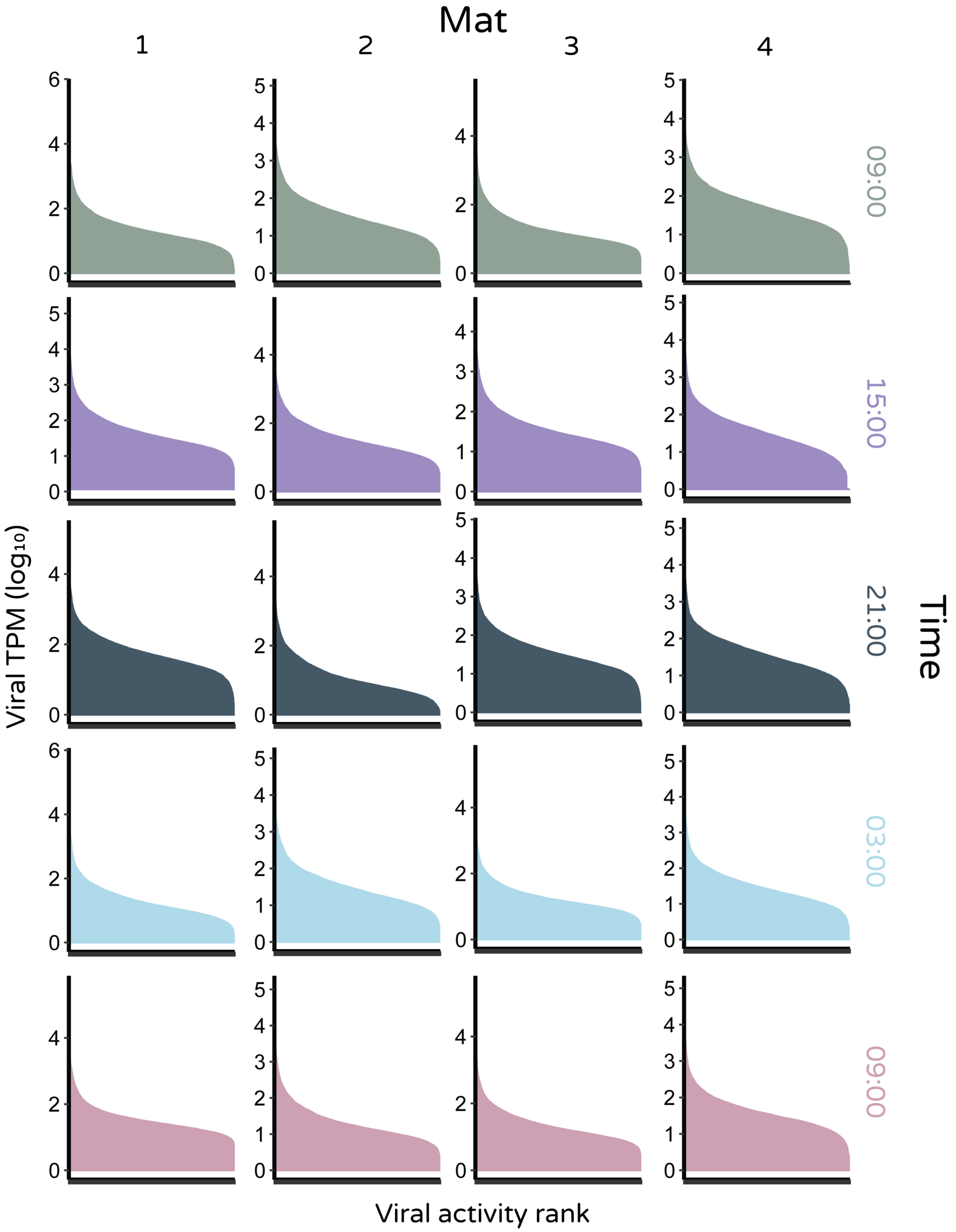


**Fig. S7.** Rank abundance ($log_{10}$TPM normalized activity [RNA]) curves across all sampled viromes (split by spatially distinct mat [x axis] and sampling time [y axis]).

**Table S1 |** Depth in meters of each mat individual sampled in this study (Mat). Also given is the relative linear distance of each mat from reference mat Individual ‘1’ on a scale of 1 to 3, with “1” the closest, and “3” the furthest (Distance). Table reproduced from Cissell and McCoy (in review).

| Mat | Depth | Distance |
|---|---|---|
| 1 | 16.4 | - |
| 2 | 16.3 | 1 |
| 3 | 16.7 | 3 |
| 4 | 16.9 | 2 |

**Table S2 |** Number of bases (Gb) and reads per sample (DNA and RNA) before (raw) and after (QC) quality trimming (Phred 25 and 100bp length for DNA; Phred 20 and 50bp length for RNA) and filtering of any human contamination (DNA only) and rRNA reads (RNA only). Reads are given as total of forward and reverse. Mean quality score across all base positions for all samples was 36. Human contaminants made up a negligible proportion of total reads (>0.01%). Percent of reads removed from quality control (including rRNA removal where appropriate), as well as GC (%) content for each sample is given. 'Total' row reports column sums or means where appropriate. Table reproduced from Cissell and McCoy (in review).

| Sample ID | Raw Bases (G) | Raw Reads (#) | QC Bases (G) | QC Reads (#) | Reads Removed (%) | GC Content (%) |
|---|---|---|---|---|---|---|
| D1_1 | 44.6 | 297 463 866 | 43.6 | 292 362 980 | 1.71 | 49 |
| D1_3 | 47.3 | 315 666 396 | 46.2 | 309 747 440 | 1.88 | 46 |
| D1_5 | 17.2 | 114 662 612 | 16.8 | 112 844 028 | 1.59 | 47 |
| D2_1 | 35.7 | 238 187 450 | 34.9 | 233 971 086 | 1.77 | 50 |
| D2_3 | 60.4 | 402 652 380 | 59.1 | 396 360 278 | 1.56 | 47 |
| D2_5 | 32.1 | 213 719 728 | 31.4 | 210 504 338 | 1.50 | 49 |
| D3_1 | 46.9 | 312 985 824 | 45.8 | 307 247 536 | 1.83 | 49 |
| D3_3 | 51.7 | 344 932 864 | 50.6 | 339 345 758 | 1.62 | 49 |
| D3_5 | 39.5 | 263 335 212 | 38.6 | 258 997 616 | 1.65 | 48 |
| D4_1 | 54.1 | 360 835 002 | 53.1 | 355 535 692 | 1.47 | 51 |
| D4_3 | 33.9 | 226 132 576 | 33.2 | 222 612 498 | 1.56 | 47 |
| D4_5 | 20.6 | 137 345 378 | 20.2 | 135 286 456 | 1.50 | 50 |
| R1_1 | 48.5 | 323 607 330 | 15.6 | 104 133 028 | 67.82 | 41 |
| R1_2 | 31.8 | 211 722 334 | 8.4 | 55 878 260 | 73.61 | 42 |
| R1_3 | 41.4 | 276 053 468 | 14.3 | 95 023 348 | 65.58 | 43 |
| R1_4 | 46.3 | 308 994 728 | 17.6 | 117 340 434 | 62.03 | 41 |
| R1_5 | 41.4 | 275 879 296 | 12.2 | 81 396 106 | 70.50 | 47 |
| R2_1 | 26.1 | 174 320 744 | 5.5 | 36 678 508 | 78.96 | 42 |
| R2_2 | 25.8 | 171 810 094 | 8.3 | 55 105 134 | 67.93 | 43 |
| R2_3 | 30.6 | 203 847 532 | 16.2 | 108 316 138 | 46.86 | 40 |
| R2_4 | 38.6 | 257 187 544 | 13.5 | 89 843 046 | 65.07 | 42 |
| R2_5 | 31.2 | 208 282 136 | 10.2 | 67 980 930 | 67.36 | 43 |
| R3_1 | 38.7 | 257 690 464 | 12.9 | 86 073 690 | 66.60 | 43 |
| R3_2 | 33.8 | 225 322 486 | 9.8 | 65 174 026 | 71.08 | 43 |
| R3_3 | 38.1 | 253 783 712 | 9.8 | 65 598 316 | 74.15 | 42 |
| R3_4 | 24.6 | 164 235 724 | 7.6 | 50 458 684 | 69.28 | 44 |
| R3_5 | 39.0 | 260 111 266 | 17.9 | 119 211 226 | 54.17 | 41 |
| R4_1 | 38.9 | 259 530 038 | 12.8 | 85 509 130 | 67.05 | 43 |
| R4_2 | 12.6 | 83 855 930 | 2.8 | 18 750 018 | 77.64 | 40 |
| R4_3 | 30.9 | 206 142 634 | 10.7 | 71 317 898 | 65.40 | 39 |
| R4_4 | 11.5 | 76 570 862 | 4.3 | 28 496 302 | 62.78 | 41 |
| R4_5 | 13.8 | 91 979 680 | 4.6 | 30 685 106 | 66.64 | 43 |
| Total | 1127.6 | 7 518 847 290 | 688.5 | 4 607 785 034 | 42.50 | 45 |

**Table S3 |** Estimates of community coverage, Nonpareil index of sequence diversity ($N_d$), and required sequencing effort to obtain 95% coverage of the underlying community derived from nonlinear regressions generated by *Nonpareil3* based on *k*-mer redundancy of *31*-mers in 10,000 randomly pulled sequences from the forward read set of each DNA sample. Rodriguez-R & Konstantinidis (2014) demonstrated that coverage values of greater than 60% perform best for assembly and detection of differential abundance. All of our samples were close to or exceeded this threshold (mean 65.46%), increasing our confidence that our taxonomic and functional inferences capture the underlying community well. 'Total' row reports column means. Table reproduced from Cissell and McCoy (in review).

| Sample | Coverage (%) | $N_d$ | Effort for 95% (bp) |
|---|---|---|---|
| D1_1 | 64.8 | 21.6 | 32 702 420 000 000 |
| D1_3 | 71.6 | 20.6 | 8 648 086 000 000 |
| D1_5 | 64.2 | 20.6 | 3 927 688 000 000 |
| D2_1 | 59.2 | 22.7 | 4 143 164 000 000 |
| D2_3 | 70.5 | 21.2 | 8 843 274 000 000 |
| D2_5 | 64.9 | 21.6 | 5 174 491 000 000 |
| D3_1 | 70.3 | 21.9 | 3 001 976 000 000 |
| D3_3 | 66.1 | 21.8 | 34 225 710 000 000 |
| D3_5 | 63.9 | 21.7 | 23 084 140 000 000 |
| D4_1 | 62.1 | 22.7 | 20 419 200 000 000 |
| D4_3 | 68.8 | 20.8 | 6 215 934 000 000 |
| D4_5 | 59.1 | 21.8 | 7 061 264 000 000 |
| Total | 65.5 | 21.6 | 13 120 600 000 000 |

**Table S4 |** Summary statistics generated using *QUAST* for contigs from DNA oassemblies (within mat across time) generated using *MEGAHIT*. Included are the total number of contigs (≥ 1500bp in length), number of contigs ≥ 50kbp in length, length of the largest contig (in bp), total assembly length (including only contigs ≥ 1500bp), N50, and L50 values. 'Total' row reports column means. Table reproduced from Cissell and McCoy (in review).

| Assembly | # Contigs ≥ 1500bp | # Contigs ≥ 50kbp | Largest Contig | Total Length | N50 | L50 |
|---|---|---|---|---|---|---|
| D1 | 305 686 | 492 | 744 424 | 956 127 986 | 3 118 | 67 787 |
| D2 | 474 013 | 584 | 1 061 358 | 1 412 520 150 | 2 894 | 114 228 |
| D3 | 512 063 | 1440 | 983 288 | 1 926 848 584 | 4 421 | 85 360 |
| D4 | 325 535 | 589 | 1 211 169 | 1 053 242 970 | 3 308 | 68 916 |
| Total | 404 324 | 776 | 1 000 060 | 1 337 184 923 | 3 435 | 84 072 |

**Table S5 |** Summary statistics generated using *METAQUAST* for contigs from cDNA (RNA) coassemblies (within mat across time) generated using *rnaSPAdes*. Included are the total number of contigs (≥ 2500bp in length), number of contigs ≥ 10kbp in length, length of the largest contig (in bp), total assembly length (including only contigs ≥ 2500bp), N50, and L50 values. 'Total' row reports column means.

| Assembly | # Contigs ≥ 2500bp | # Contigs ≥ 10kbp | Largest Contig | Total Length | N50 | L50 |
|---|---|---|---|---|---|---|
| R1 | 41 361 | 795 | 93 571 | 46 771 946 | 5 845 | 2 305 |
| R2 | 29 925 | 560 | 62 183 | 37 626 848 | 5 530 | 2 063 |
| R3 | 38 824 | 959 | 56 831 | 41 463 656 | 7 375 | 1 542 |
| R4 | 19 462 | 174 | 84 790 | 14 978 477 | 5 144 | 923 |
| Total | 32 393 | 622 | 74 344 | 35 210 232 | 5 974 | 1 708 |
